# Lack of universal mutational biases in a fungal phylum

**DOI:** 10.1101/2022.03.29.486229

**Authors:** Qianhui Zheng, Jacob L. Steenwyk, Antonis Rokas

## Abstract

Mutations fuel the diversity of life forms on earth through changes of nucleotides in DNA sequence. Patterns of mutations are biased; for example, mutational biases toward adenine and thymine have been previously noted in bacteria and biases for transitions over transversions are observed in diverse groups of organisms. However, the mutational biases in fungi, whose genomes vary widely in their GC content, remain poorly understood. Here, we characterized patterns of single nucleotide polymorphisms among 537 strains from 30 species and four classes from Ascomycota, the most species-rich fungal phylum. We found that mutational biases vary across Ascomycota; for example, some species in the class Saccharomycetes, in particular the bipolar budding yeast *Hanseniaspora uvarum* and the emerging pathogen *Candida auris*, show strong mutational bias toward A|T substitutions whereas the black mold *Stachybotrys chartarum* in the class Sordariomycetes shows substantial mutational bias toward G|C substitutions. Examination of GC content and GC equilibrium content, a measure that represents the GC content under selective neutrality and accounts for rates of G|C > A|T and A|T > G|C substitutions, revealed that fungal species vary in how their genome nucleotide composition is affected by neutral processes, mutational biases, and external evolutionary forces, such as selection. Whereas genome nucleotide composition is consistent with neutral expectations and is mainly driven by mutational bias in some species (e.g., *Zymoseptoria tritici*), the composition of others is influenced by both mutational bias and selection (e.g., *H. uvarum* and *S. chartarum*). These results highlight the variation of patterns of mutations across a fungal phylum and suggest that both neutral and selective processes shape the nucleotide composition of fungal genomes.

## Introduction

Mutations are the underlying force for genetic novelty and variability (Barton 2010), serving as the raw material for evolutionary change. Comprehensive knowledge of patterns of mutations is critical for understanding the molecular mechanisms of evolution. Although the randomness of mutations and what we precisely mean by “randomness” continues to be debated (Hershberg and Petrov 2010; Monroe et al. 2022; Eyre-Walker and Keightley 2007; Haag-Liautard et al. 2007; Orr 2005), it is well accepted that various factors, such as the different biochemical properties of nucleotides, make certain mutations more likely to occur than others resulting in unbalanced or biased rates of mutations. Detailed investigation of mutational biases can shed light on variation in genome nucleotide composition and help refine our understanding of the molecular mechanisms that contribute to evolution.

Previous studies suggest that mutations may be universally biased toward the A|T direction in diverse organisms. For example, Hershberg and Petrov showed that mutations are biased toward A|T in bacteria whose genomes varied widely in their GC contents, suggesting that bacterial mutational biases are far less variable than previously thought (Hershberg and Petrov 2010). Similarly, Keightley et al. observed that the G|C > A|T mutation rate is close to two times that of the A|T > G|C mutation rate in three *Drosophila melanogaster* mutation accumulation lines (Keightley et al. 2009), and Lynch used data on *de novo* mutations for monogenic disorders to show a preponderance of mutations toward A|T in the human genome (Lynch 2010). In all these studies, the disparity between the observed GC content and the neutral expectation that A|T-biased mutations would lead to an AT-rich genome suggests that genome nucleotide composition is not at an equilibrium determined by mutational biases alone. Countering forces such as natural selection and biased gene conversion (BGC) must be acting in favor of G|C mutations to achieve the observed GC content, particularly for species with GC-rich genomes (Hershberg and Petrov 2010; Webster, Smith, and Ellegren 2003; Haddrill and Charlesworth 2008).

In contrast to other major lineages—such as bacteria, animals, and plants—the mutational biases of organisms from the fungal kingdom remain severely understudied. Some studies suggest that fungi, much like bacteria and animals, show a mutational bias toward A|T. For example, Zhu et al. observed a strong A|T bias of mutations driven by both C>T transitions and C>A transversions in mutation accumulation lines of the budding yeast *Saccharomyces cerevisiae* (Zhu et al. 2014). An excess of C>T mutations can also stem from processes such as repeat-induced point mutation (RIP), a defense mechanism against invasive transposable elements in *Neurospora crassa* (Selker et al. 1987; Cambareri et al. 1989) and other fungal species (Galagan and Selker 2004; Horns et al. 2012; Hane et al. 2014). Although these insights are valuable, they are based on studies of a few species; whether mutations are biased toward the A|T direction across fungi in general remains an open question.

Species in the fungal kingdom substantially vary in genome composition and content, morphologies, and ecological functions (Li et al. 2021). It is estimated that fungi comprise between 2.2 to 3.8 million species, but only around 150,000 have been formally described (Hawksworth and Lücking 2017; Willis 2018). Ascomycota contains three subphyla—Pezizomycotina, Saccharomycotina, and Taphrinomycotina—and is the largest phylum of fungi representing around two□thirds of all described species (Shen et al. 2020; James et al. 2020; Naranjo-Ortiz and Gabaldón 2019). Draft genomes for more than 1,000 Ascomycota species are publicly available, making the group one of the most sequenced phyla (Shen et al. 2020). Ascomycetes exhibit a variety of lifestyles, such as yeasts, pathogens, parasites, saprobes, endophytes, and molds (Naranjo-Ortiz and Gabaldón 2019). Both the biological diversity and the amount of available genome data make Ascomycota highly suitable for studying mutational bias and its impact on genome sequence evolution.

In this study, we examined mutational biases in 537 strains from 30 diverse species from four different classes of the phylum Ascomycota. To minimize the influence of selection and BGC on our results, mutational patterns were measured from single nucleotide polymorphism (SNP) data reasoning that selection and BGC will likely be less effective on sites that are polymorphic within species than on sites that have diverged between species (Akashi 1995; Messer 2009). The resulting SNPs were used to calculate relative genome-wide rates of nucleotide mutations, transition over transversion ratios, directional mutational biases, and GC equilibrium contents. We find that mutational biases are highly variable across species in Ascomycota. While species in the class Saccharomycetes, in particular *Hanseniaspora uvarum* and *Candida auris*, show A|T mutational bias, mutations are biased toward G|C in *Stachybotrys chartarum* (class Sordariomycetes) and neutral in *Aspergillus fumigatus* (class Eurotiomycetes). Examination of GC equilibrium and observed GC contents revealed that evolutionary forces (e.g., selection) do not uniformly affect genome nucleotide compositions of species in Ascomycota. Evolutionary forces could be G|C biased (*H. uvarum*), A|T biased (*S. chartarum*), or neutral (*Z. tritici*). As a result, selection may magnify the direction of mutational bias, as in the case of *Purpureocillium lilacinum*, or counterbalance it, as in the case of *S. chartarum*. Although previous studies in many organisms have shown that mutational bias is generally toward A|T with selection countering it by favoring mutations toward G|C, our study suggests that mutational biases and selection vary in both their direction and strength in fungal genomes.

## Materials and Methods

### Data Set Assembly and Curation

We chose 30 species of Ascomycota based on their broad phylogenetic distribution and the fact that each species has at least three strains with a publicly available genome assembly. The sister species to each of our 30 chosen species were determined according to a recently published genome-scale phylogeny of 1,107 species from Ascomycota (Shen et al. 2020). All genome assemblies were downloaded from NCBI (https://www.ncbi.nlm.nih.gov/). Genome statistics including genome size, N50 values, and number of scaffolds that provide an overview of the quality of the genomes of all strains used in this study are presented in Table S1.

To ensure that strains included in our analyses were not taxonomically misclassified or represent organisms from a complex comprised of multiple species (Jos et al. 2022), we calculated the pairwise intra-species average nucleotide identity (ANI) values for all strains within a given species using FastANI (https://github.com/ParBLiSS/FastANI) (Figure S1). We used 90% ANI as the cutoff for deciding whether a strain should belong to a species. Using this cutoff, three isolates—*Aureobasidium pullulans* AY4 (ANI value: 80.75±1.05), *Aureobasidium pullulans* CCTCC-M-2012259 (ANI value: 80.79±1.04), *Fusarium fujikuroi* FUS01 (although the ANI value 90.27±0.10 is above 90, it still largely deviates from the other strains of the species), and *Yarrowia lipolytica* NCIM3590 (ANI value: 80.73±0.02)—were misclassified and removed from our analyses. ANI analysis of the genomes *Penicillium expansum* YT02 and *Aspergillus fumigatus* SPS-2 strains failed to complete. Examination of GC contents, transition over transversion ratios, and directional mutational biases in these two strains revealed they are outliers, a finding suggesting that they may be misclassified; thus, we erred on the side of caution and removed them from our analyses. After removing these isolates, the final data set is comprised of 537 strains from 30 species (Table S1). A full list of strains and species analyzed in this study, along with the corresponding sister species and genome statistics, is presented in Table 1.

**Table 1.** Summary of the 30 species analyzed and their genome statistics.

| Group | Number of<br>genomes<br>(strains)<br>analyzed | Mean ANI | Outgroup | Mean SNPs |
| --- | --- | --- | --- | --- |
| **Dothideomycetes** | **98** |  |  | **999789 (s = 496652)** |
| *Alternaria alternata* | 11 | 98.30 | *Alternaria arborescens* | 882832 (s = 10395) |
| *Alternaria arborescens* | 6 | 99.33 | *Alternaria alternata* | 881431 (s = 2641) |
| *Aureobasidium pullulans* | 52 | 97.46 | *Aureobasidium melanogenum* | 632417 (s = 46396) |
| *Pyrenophora tritici-repentis* | 10 | 99.84 | *Pyrenophora teres* | 2008903 (s = 7294) |
| *Zymoseptoria tritici* | 19 | 98.04 | *Zymoseptoria brevis* | 1579202 (s = 5794) |
| **Eurotiomycetes** | **175** |  |  | **1269157 (s = 605868)** |
| *Aspergillus flavus* | 104 | 98.97 | *Aspergillus arachidicola* | 1633936 (s = 15750) |
| *Aspergillus fumigatus* | 23 | 99.57 | *Aspergillus fischeri* | 1565477 (s = 5864) |
| *Blastomyces dermatitidis* | 3 | 98.52 | *Blastomyces gilchristii* | 74139 (s = 60423) |
| *Coccidioides immitis* | 5 | 99.24 | *Coccidioides posadasii* | 354803 (s = 5444) |
| *Coccidioides posadasii* | 10 | 98.80 | *Coccidioides immitis* | 340341 (s = 11946) |
| *Histoplasma capsulatum* | 7 | 96.07 | *Blastomyces gilchristii* | 838338 (s = 41090) |
| *Paracoccidioides brasiliensis* | 4 | 99.07 | *Paracoccidioides lutzii* | 1112522 (s = 9715) |
| *Penicillium expansum* | 7 | 99.77 | *Penicillium oxalicum* | 40821 (s = 2649) |
| *Trichophyton rubrum* | 12 | 99.93 | *Trichophyton soudanense* | 13595 (s = 683) |
| **Sordariomycetes** | **103** |  |  | **732059 (s = 679956)** |
| *Beauveria bassiana* | 12 | 97.50 | *Beauveria rudraprayagi* | 395456 (s = 309975) |
| *Colletotrichum higginsianum* | 3 | 99.71 | *Colletotrichum lentis* | 2313553 (s = 139100) |
| *Fusarium fujikuroi* | 17 | 99.36 | *Fusarium proliferatum* | 1128606 (s = 29714) |
| *Fusarium proliferatum* | 13 | 98.71 | *Fusarium fujikuroi* | 1093463 (s = 11969) |
| *Metarhizium anisopliae* | 6 | 98.05 | *Metarhizium robertsii* | 840379 (s = 52866) |
| *Purpureocillium lilacinum* | 6 | 98.85 | *Drechmeria coniospora* | 209694 (s = 5364) |
| *Stachybotrys chartarum* | 4 | 99.25 | *Stachybotrys chlorohalonata* | 1933190 (s = 26173) |
| *Trichoderma harzianum* | 6 | 94.95 | *Trichoderma virens* | 2091776 (s = 39920) |
| *Verticillium dahliae* | 36 | 99.31 | *Verticillium longisporum* | 103631 (s = 2590) |
| **Saccharomycetes** | **161** |  |  | **514171 (s = 376637)** |
| *Candida albicans* | 49 | 99.12 | *Candida dubliniensis* | 832895 (s = 171050) |
| *Candida auris* | 58 | 99.70 | *Candida heveicola* | 94437 (s = 2504) |
| *Hanseniaspora uvarum* | 10 | 98.76 | *Hanseniaspora nectarophila* | 293501 (s = 22810) |
| *Komagataella phaffii* | 6 | 99.99 | *Komagataella populi* | 607399 (s = 1484) |
| *Saccharomyces paradoxus* | 13 | 97.22 | *Saccharomyces cerevisiae* | 1092589 (s = 11495) |
| *Yarrowia lipolytica* | 22 | 99.69 | *Yarrowia keelungensis* | 697308 (s = 10992) |
| *Zygosaccharomyces bailii* | 3 | 99.55 | *Zygosaccharomyces bisporus* | 122833 (s = 29210) |

### Identification of Single Nucleotide Polymorphisms

To identify and polarize the direction of mutations in SNPs within each species, we used MUMmer 3.0 (Kurtz et al. 2004), a toolkit for whole genome comparisons, as our main analytical tool (http://mummer.sourceforge.net/). To do so, we first ran the NUCmer function (NUCleotide MUMmer) included in MUMmer to align genome assemblies. This required two inputs: a reference genome and a query genome to identify SNPs in. For the reference genome, the most closely related species of the query species, which we identified from a recently published genome-scale phylogeny of 1,107 Ascomycota genomes (Shen et al. 2020), was used. This analysis allowed us to polarize the direction of mutations in SNPs following a previously established protocol (Hershberg and Petrov 2010). The resulting NUCmer output was used as input to the show-snps function in MUMmer. To ensure SNPs were robustly identified, ambiguously mapped alignments were removed.

### Estimation of Mutational Bias

The resulting SNPs were used to examine mutational biases. To determine the relative rates of the six pairwise substitutions, we determined the number of SNPs that exhibited guanine (G)-to-adenine (A) and cytosine (C)-to-thymine (T) (G>A|C>T), A>G|T>C, A>C|T>G, G>T|C>A, A>T|T>A, and G>C|C>G mutations across both protein-coding regions and non-coding regions of genomes. To determine whether a variant occurred in protein-coding regions, SNP positions were cross-referenced with the predicted gene boundaries (available via gtf/gff files) of the sister species when gene annotations were available. Files detailing gene boundaries were downloaded from NCBI. Eight of the 30 species examined lacked publicly available gene annotations, so our examination of protein-coding and non-coding patterns of SNPs was performed in the remaining 22 genomes. We also calculated the transition-to-transversions rate (Ts/Tv) and the number of variants that have a G|C in the sister species and A|T in the query species over the number of variants that have A|T in the sister species and G|C in the query species.

To evaluate directional biases in the observed patterns of mutations, the observed GC content and the expected equilibrium GC content for each species and class was calculated. GC_eq_ is calculated from the measured rates of A|T > G|C and G|C > A|T mutations using the following equation (Hershberg and Petrov 2010):

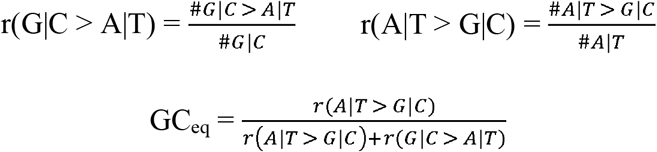

The aforementioned analyses were conducted using custom scripts written in the Python programming language (https://www.python.org/), which are available via figshare (see Data Availability Statement). The resulting data were summarized using ggplot2, v3.3.5 (https://www.rdocumentation.org/packages/ggplot2/) and statistical analyses of resulting data were conducted using the R, v3.6.3 (https://cran.r-project.org/), programming language.

### Data Availability Statement

All data and custom scripts necessary for replicating the results of this study will be freely available upon publication in figshare.

## Results

### The genomes of species in Ascomycota vary widely in their GC contents

By calculating average GC content for each species and each class (Figure 1 and Table 2), we found that species in Ascomycota vary widely in their GC contents. The average GC content per class is 51.23 ± 0.84% for Dothideomycetes, 48.55 ± 1.28 % for Eurotiomycetes, 52.01 ± 2.52% for Sordariomycetes, and 42.07 ± 5.07% for Saccharomycetes. Thus, Saccharomycetes have considerably lower GC content in comparison to the other three classes. Among the thirty species, *Purpureocillium lilacinum* (Sordariomycetes) has the highest GC content 57.29 ± 0.26% and *Hanseniaspora uvarum* (Saccharomycetes) has the lowest GC content 34.53 ± 0.29%.

**Table 2.** GC_eq_ values, GC content, and their correlation across fungal classes and species.

| **Group** | **GC<sub>eq</sub>** | **GC**<br>**content** | **$\Delta_{GC - GC_{eq}}$** | **Correlation coefficient**<br>**(p-value)** |
| --- | --- | --- | --- | --- |
| **Dothideomycetes** | **0.53** | **0.51** | **-0.02** | **0.04 (p = 6.95E-1)** |
| *Alternaria alternata* | 0.53 | 0.51 | -0.02 | 0.93 (p = 2.50E-5) |
| *Alternaria arborescens* | 0.49 | 0.51 | 0.02 | 0.88 (p = 2.06E-2) |
| *Aureobasidium pullulans* | 0.54 | 0.51 | -0.03 | 0.92 (p = 3.67E-22) |
| *Pyrenophora tritici-repentis* | 0.50 | 0.51 | 0.01 | 0.99 (p = 4.73E-8) |
| *Zymoseptoria tritici* | 0.53 | 0.53 | 0 | 0.99 (p = 6.05E-16) |
| **Eurotiomycetes** | **0.48** | **0.49** | **0.01** | **0.83 (p = 8.08E-46)** |
| *Aspergillus flavus* | 0.49 | 0.49 | 0 | 0.35 (p = 2.69E-4) |
| *Aspergillus fumigatus* | 0.50 | 0.50 | 0 | 0.96 (p = 3.79E-13) |
| *Blastomyces dermatitidis* | 0.43 | 0.44 | 0.01 | 0.40 (p = 7.39E-1) |
| *Coccidioides immitis* | 0.49 | 0.48 | -0.01 | 0.91 (p = 2.99E-2) |
| *Coccidioides posadasii* | 0.48 | 0.48 | 0 | 0.67 (p = 3.31E-2) |
| *Histoplasma capsulatum* | 0.42 | 0.44 | 0.02 | 1.00 (p = 2.21E-10) |
| *Paracoccidioides brasiliensis* | 0.50 | 0.47 | -0.03 | 0.47 (p = 5.27E-1) |
| *Penicillium expansum* | 0.47 | 0.49 | 0.02 | 0.59 (p = 1.65E-1) |
| *Trichophyton rubrum* | 0.49 | 0.49 | 0 | 0.81 (p = 1.29E-3) |
| **Sordariomycetes** | **0.52** | **0.52** | **0** | **0.77 (p = 4.94E-21)** |
| *Beauveria bassiana* | 0.52 | 0.51 | -0.01 | 0.15 (p = 6.35E-1) |
| *Colletotrichum higginsianum* | 0.50 | 0.54 | 0.04 | 1.00 (p = 4.71E-2) |
| *Fusarium fujikuroi* | 0.49 | 0.49 | 0 | 0.83 (p = 4.57E-5) |
| *Fusarium proliferatum* | 0.49 | 0.49 | 0 | 0.97 (p = 4.69E-8) |
| *Metarhizium anisopliae* | 0.52 | 0.52 | 0 | 0.43 (p = 3.91E-1) |
| *Purpureocillium lilacinum* | 0.54 | 0.57 | 0.03 | 0.96 (p = 2.69E-3) |
| *Stachybotrys chartarum* | 0.58 | 0.54 | -0.04 | 0.96 (p = 3.54E-2) |
| *Trichoderma harzianum* | 0.51 | 0.50 | -0.01 | 1.00 (p = 1.18E-5) |
| *Verticillium dahliae* | 0.54 | 0.54 | 0 | 0.77 (p = 3.58E-8) |
| **Saccharomycetes** | **0.37** | **0.42** | **0.05** | **0.52 (p = 1.92E-12)** |
| *Candida albicans* | 0.37 | 0.36 | -0.01 | 0.78 (p = 4.76E-11) |
| *Candida auris* | 0.36 | 0.46 | 0.1 | 0.82 (p = 4.11E-15) |
| *Hanseniaspora uvarum* | 0.24 | 0.35 | 0.11 | 0.97 (p = 1.81E-6) |
| *Komagataella phaffii* | 0.40 | 0.42 | 0.02 | 1.00 (p = 2.94E-6) |
| *Saccharomyces paradoxus* | 0.41 | 0.40 | -0.01 | 0.85 (p = 2.27E-4) |
| *Yarrowia lipolytica* | 0.43 | 0.49 | 0.06 | 0.92 (p = 7.79E-10) |
| *Zygosaccharomyces bailii* | 0.38 | 0.44 | 0.06 | 0.72 (p = 4.92E-1) |

**Figure 1.**
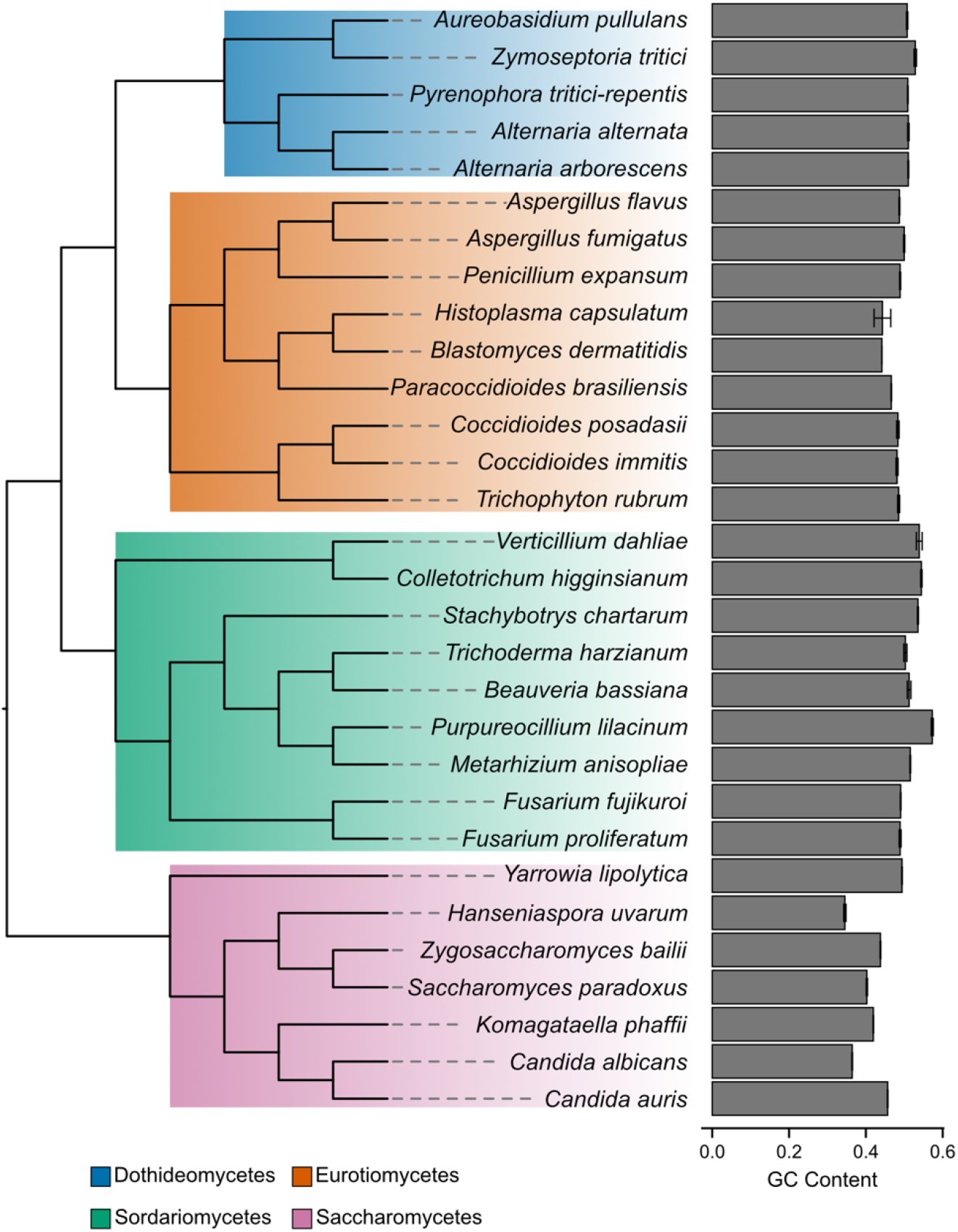
Species from the fungal phylum Ascomycota vary in their GC contents. The phylogeny of the 30 species examined in this study, along with their observed genome-wide GC contents (as a fraction). The 30 species belong to 4 classes: Dothideomycetes (5 species; 100 strains), Eurotiomycetes (9 species; 167 strains), Sordariomycetes (9 species; 91 strains), and Saccharomycetes (7 species; 160 strains). Most of the species have GC contents around 50%. The average GC content per class is 51.23 ± 0.84% for Dothideomycetes, 48.55 ± 1.28 % for Eurotiomycetes, 52.01 ± 2.52% for Sordariomycetes, and 42.07 ± 5.07% for Saccharomycetes. *Purpureocillium lilacinum* has the highest GC content with 57.29 ± 0.26% and *Hanseniaspora uvarum* has the lowest GC content with 34.53 ± 0.29%. Detailed GC contents for all species are summarized in Table 2. The phylogeny is based off evolutionary relationships from a genome-scale phylogeny of 1,107 species from Ascomycota (Shen et al. 2020); the subtree of the 30 species examined in this study was obtained using treehouse (Steenwyk and Rokas 2019).

### *Purpureocillium lilacinum* exhibits an unusually low transition to transversion ratio

Examination of transitions and transversions showed that transitions typically occur twice as frequently as transversions across all species apart from *P. lilacinum*, which has exceptionally high G>C|C>G transversion rate and a transition over transversion ratio of 0.79±0.00 (Figure 2, Figure S3). Since genome composition could affect the relative rates of different categories of SNPs, the high G>C|C>G transversion rate of *P. lilacinum* may be explained by its particularly high GC content. Likewise, *Candida albicans* and *H. uvarum*, which have the lowest GC contents, show comparably higher A>T|T>A transversion rates. Further examination of protein-coding and non-coding patterns of SNPs reveals that these mutational biases operate consistently across protein-coding and non-coding regions of genomes (Figure S2).

**Figure 2.**
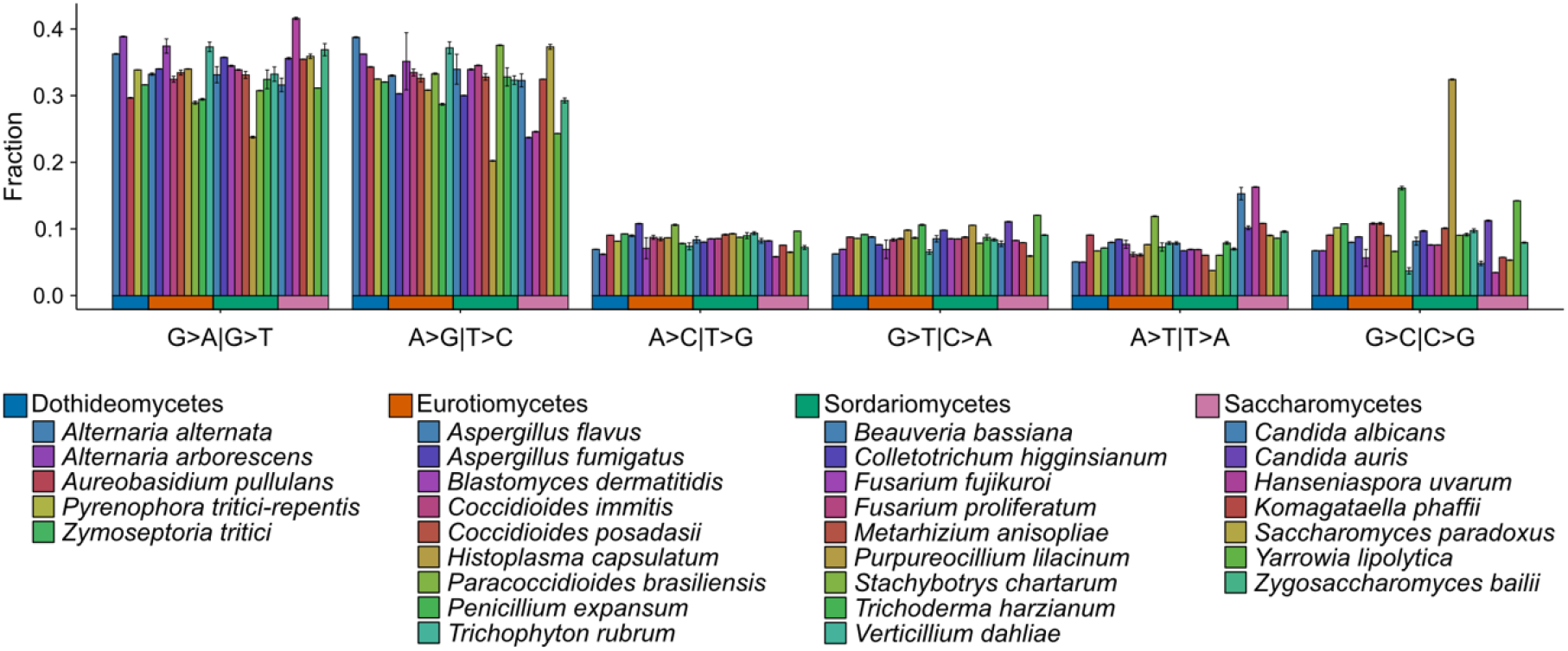
Relative genome-wide rates of different types of nucleotide mutations across Ascomycota fungi. Transitions (G>A|C>T and A>G|T>C) occurred more frequently than transversions in 29 of the 30 species’ genomes examined. The sole exception was *Purpureocillium lilacinum*, a species with high G>C|C>G transversion rate.

### Saccharomycetes species show strong A|T mutational biases while *Stachybotrys chartarum* (Sordariomycetes) shows G|C mutational bias

To measure directional mutational bias, we calculated the ratio of G|C > A|T substitutions over A|T > G|C substitutions (Figure 3). We found that most species in Dothideomycetes, Eurotiomycetes, and Sordariomycetes have G|C > A|T over A|T > G|C ratios close to 1, suggesting that there is no substantial mutational bias. In contrast, some species in Saccharomycetes, in particular *Candida auris* and *H. uvarum*, demonstrate strong mutational bias toward A|T, with their G|C > A|T over A|T > G|C ratios being 1.46 ± 0.01 and 1.64 ± 0.01, respectively. Since this measure does not consider GC content, the G|C > A|T over A|T > G|C ratio could appear higher for species with higher GC content, as the cases of *Colletotrichum higginsianum* and *P. lilacinum*. We also note that *Stachybotrys chartarum* has the lowest G|C > A|T over A|T > G|C ratio at 0.83 ± 0.00, which is suggestive of substantial mutational bias toward G|C. These results suggest that there is no universal pattern of mutational bias in Ascomycota.

**Figure 3.**
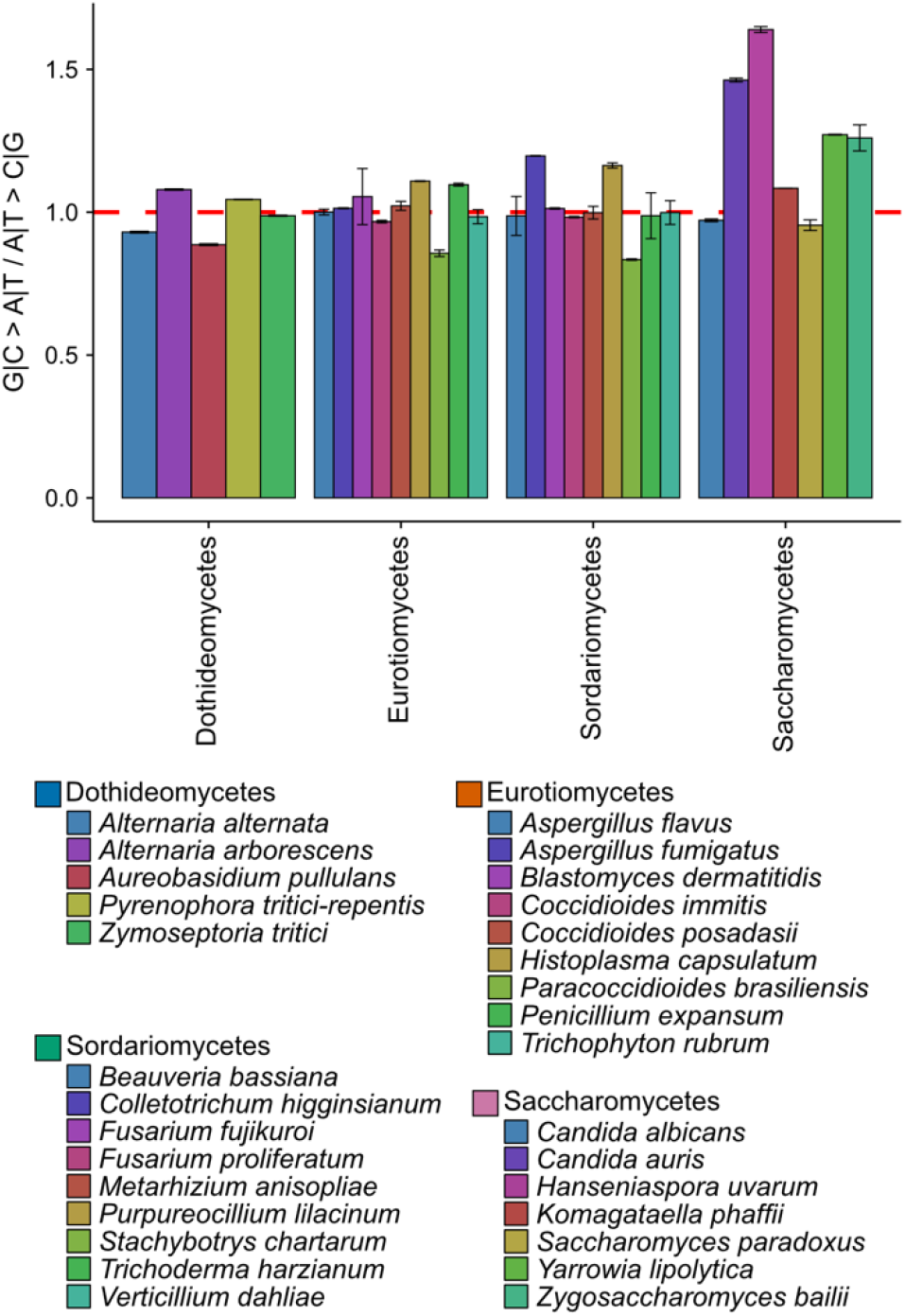
Species from the class Saccharomycetes exhibit the strongest mutational bias toward A|T. Examination of G|C > A|T substitutions over A|T > G|C substitutions, a measure of directional mutation bias, revealed that most species in Dothideomycetes, Eurotiomycetes, and Sordariomycetes have G|C > A|T over A|T > G|C ratios around one (red dashed line), suggesting the absence of substantial mutational bias. In contrast, several species in Saccharomycetes (budding yeasts), most notably as the emerging human pathogen *Candida auris* and the bipolar yeast *Hanseniaspora uvarum*, exhibit strong mutational bias toward A|T.

To further understand the direction of mutational bias in fungi, we calculated GC equilibrium content, which represents the expected GC content under the observed rates of G|C > A|T and A|T > G|C substitutions (Hershberg and Petrov 2010). If mutational bias is neutral, then the rate of A|T > G|C substitutions should equal the rate of G|C > A|T substitutions and GC_eq_ should be 0.50. If mutations are biased toward A|T, then the rate of the rate of G|C > A|T substitutions should exceed the rate of A|T > G|C substitutions and GC_eq_ should be smaller than 0.50. Conversely if mutations are biased toward G|C, then the rate of the rate of A|T > G|C substitutions should exceed the rate of G|C > A|T substitutions and GC_eq_ should be greater than 0.50. Examination of GC_eq_ values shows that species in Ascomycota vary widely, suggestive of diverse mutational biases (Table 2). The average GC_eq_ per class is 0.53 ± 0.02 for Dothideomycetes, 0.48 ± 0.02 for Eurotiomycetes, 0.52 ± 0.03 for Sordariomycetes, and 0.37 ± 0.04 for Saccharomycetes. The strongest A|T bias was observed in *C. auris* and *H. uvarum*, which have GC_eq_ values of 0.37 ± 0.00 and 0.24 ± 0.00, respectively. A substantial G|C mutational bias was observed in *S. chartarum* (GC_eq_ = 0.58 ± 0.00).

### Differences in GC content across Ascomycota fungi can be shaped by selection

GC_eq_ should approximate current GC content if genome nucleotide composition is evolving neutrally and is solely influenced by mutational bias. In contrast, a significant difference between GC_eq_ and current GC content would suggest that genome nucleotide composition is also influenced by other evolutionary forces, such as natural selection. Thus, by comparing GC_eq_ value and observed GC content for each species in our study, the impact (and strength) of selection on genome nucleotide composition can be examined. To do so, we plotted GC_eq_ against GC content for each species and each class as well as their regression lines using a linear model and calculated the Δ_GC–GC_eq__ value, a measure of the direction and strength of natural selection (Figure 4; Table 2). For species whose Δ_GC–GC_eq__ values are around 0, genome nucleotide composition is mostly determined by mutational biases. For species whose Δ_GC–GC_eq__ values are positive, genome nucleotide composition is determined by mutational biases and selection that favors G|C nucleotides; for species whose Δ_GC–GC_eq__ values are negative, genome nucleotide composition is determined by mutational biases and selection that favors A|T nucleotides.

**Figure 4.**
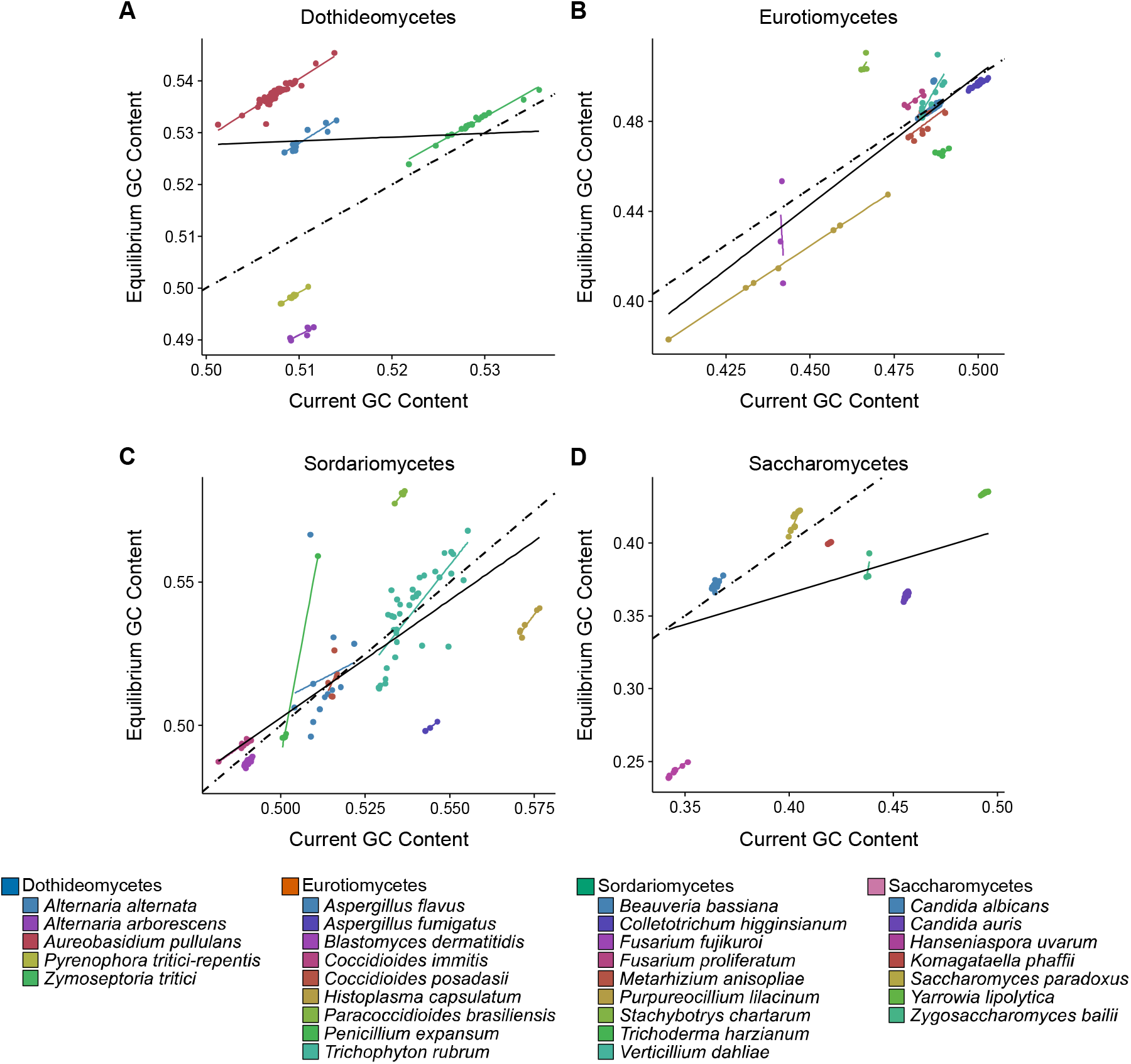
Variation in mutational biases between and within species of classes of Ascomycota. The relationship between equilibrium GC content (GC_eq_) and the observed GC content values for species in Dothideomycetes (A), Eurotiomycetes (B), Sordariomycetes (C), and Saccharomycetes (D). GC_eq_ represents the expected GC content with observed rates of G|C > A|T and A|T > G|C substitutions. Each colored dot corresponds to the genome of a strain from a given species and each colored line represents the least squares regression line for one species; each solid black line represents the regression line for one class; each dashed black line represents the relationship between GC_eq_ and the current GC content values when they are equal. Note that *H. uvarum* (Saccharomycetes) and *S. chartarum* (Sordariomycetes) deviate most from the equality line. Detailed class- and species-specific GC_eq_, GC content, and the correlation coefficient values are summarized in Table 2. Species colors are the same as in Figure 2.

For most species in Eurotiomycetes and Sordariomycetes, the GC_eq_ and GC content values are roughly equal, suggesting that nucleotide composition of these genomes is primarily, if not exclusively, shaped by mutational bias. This is the case both for species with mutational biases toward A|T (e.g., *Coccidioides posadasii*; GC_eq_ = 0.48; GC = 0.48) and species with mutational biases toward G|C (e.g., *Verticillium dahliae*; GC_eq_ = 0.54; GC = 0.54). Two notable exceptions include two Sordariomycetes: *P. lilacinum* (GC_eq_ = 0.54; GC = 0.57) and *S. chartarum* (GC_eq_ = 0.58; GC = 0.54). GC_eq_ is lower than GC content in *P. lilacinum*, suggesting that its genome composition is influenced by mutations and selection that are both G|C-biased, whereas GC_eq_ is higher than GC content in *S. chartarum*, suggesting that its genome composition is influenced by G|C-biased mutations and A|T-biased selection.

In Dothideomycetes and Saccharomycetes there are more species whose GC_eq_ and GC content values deviate from a 1:1 ratio meaning that their genome compositions are shaped by both mutational bias and selection. In Dothideomycetes, *Alternaria alternata* (GC_eq_ = 0.53; GC = 0.51) and *Aureobasidium pullulans* (GC_eq_ = 0.54; GC = 0.51) show evidence of A|T-biased selection. In Saccharomycetes, selection is generally biased toward G|C, possibly to counterbalance the A|T-biased mutations. The effects of G|C-biased selection are most notable in *H. uvarum* (GC_eq_ = 0.24; GC = 0.35) and *C. auris* (GC_eq_ = 0.36; GC = 0.46), who possess the strongest mutational biases toward A|T. However, some species such as *Zymoseptoria tritici* (GC_eq_ = 0.53; GC = 0.53) in Dothideomycetes and *C. albicans* (GC_eq_ = 0.37; GC = 0.36) in Saccharomycetes do not appear to be influenced by selection.

## Discussion

In this study, we analyzed patterns of mutations in SNPs from population genomic data in 30 Ascomycota species. In contrast with expectations based on previous studies that mutations are universally biased toward A|T, our results suggest that species in Ascomycota exhibit diverse mutational biases. While most species lack substantial mutational bias, mutations are evidently biased toward A|T in Saccharomycetes, particularly in *H. uvarum and C. auris*. Furthermore, evolutionary forces such as selection do not always favor G|C to counteract mutational biases, as observed in previous studies. Selection varies in both direction and strength across different species.

### Species in Ascomycota exhibit substantial diversity in their mutational biases

It is well established that some nucleotides are more unstable than others giving rise to biases in the frequencies of mutations in genomes. For example, deamination of methylated cytosines to thymine occurs at high frequency, contributing to A|T bias in spontaneous mutations. Indeed, several previous studies found that mutations are biased toward A|T in many organisms (Hershberg and Petrov 2010; Keightley et al. 2009; Lynch 2010; Webster, Smith, and Ellegren 2003), raising the hypothesis that mutational bias toward A|T may be universal. However, our examination of 537 strains from 30 fungal species in the phylum Ascomycota revealed that this is not the case. Rather, we found that the mutational biases of these species are quite diverse (Figure 3; Table 2). For example, *H. uvarum* and *C. auris* in the class Saccharomycetes have pronounced A|T mutational bias, whereas *S. chartarum* in the class Sordariomycetes exhibits considerable G|C mutational bias. There are also several species that appear to lack mutational bias, such as *Aspergillus fumigatus* in the class Eurotiomycetes. This diversity of mutational biases is not only observed between fungal classes but is also observed when we compare patterns of mutations within each class. For example, within the class Dothideomycetes, the species *Alternaria alternata*, *Alternaria arborescens*, *Aureobasidium pullulans*, and *Pyrenophora tritici-repentis* all have GC content values around 0.51 but their mutational biases may be A|T biased, G|C biased, or lack evidence of bias (Figure 3; Table 2).

In general, Saccharomycetes have lower GC content and stronger A|T mutational bias than the other three classes. A previous comparison of the subphylum Saccharomycotina (Saccharomycetes is the only class within the subphylum) to Pezizomycotina (a subphylum that contains more than a dozen classes, including Dothideomycetes, Eurotiomycetes, and Sordariomycetes) showed that Saccharomycotina exhibit higher evolutionary rates than Pezizomycotina and contain smaller sets of DNA repair genes (Shen et al. 2020). Thus, the pronounced mutational bias toward A|T may be associated with the lower number of DNA repair genes in Saccharomycetes. Furthermore, Shen et al. found that evolutionary rate shows negative correlation with GC content in Saccharomycotina but positive correlation in Pezizomycotina (Shen et al. 2020). Since our results suggest that Saccharomycotina have strong mutational biases toward A|T but Pezizomycotina mostly lack or have weak mutational biases toward G|C, evolutionary rate may be associated with the strength of mutational bias across the entire Ascomycota phylum.

### The A|T mutational bias of H. uvarum is likely related to its extensive loss of DNA repair genes

The bipolar budding yeast *H. uvarum* is commonly found on grapes and is often very abundant at the beginning of alcoholic fermentation (Albertin et al. 2016; Schütz and Gafner 2008; Hierro et al. 2006). Steenwyk et al. previously showed that *H. uvarum* belongs to a faster-evolving lineage (FEL) of the genus *Hanseniaspora* (Steenwyk et al. 2019). The FEL stem branch experienced accelerated evolutionary rate, which coincides with an extensive loss of genes in metabolic, cell cycle regulation, and DNA repair pathways (Steenwyk et al. 2019). The FEL also has strong mutational signatures associated with 8-oxo-deoxyguanosine and UV damage (Steenwyk et al. 2019), which result in G|C>A|T mutations. Our observation of the A|T mutational bias in *H. uvarum* is consistent with Steenwyk et al.’s finding that substitutions are biased towards A|T in the FEL. The A|T mutational bias of *H. uvarum* may in part be explained by DNA damage by diverse mutagens including endogenous mutagens such as 8□oxo□deoxyguanosine and exogenous mutagens such as UV radiation (de Bont and van Larebeke 2004; Budden and Bowden 2013). The example of *H. uvarum* provides additional support for the potential association between A|T mutational bias and loss of DNA repair genes.

### The A|T mutational bias of C. auris may be linked to its ability to quickly evolve multidrug resistance

*C. auris* is a newly emerging multidrug-resistant fungal pathogen. It was first described in 2009 in Japan (Satoh et al. 2009), and has caused nosocomial outbreaks on five continents (Jeffery-Smith et al. 2017). Due to its rapid spread in hospitalized patients and the difficulties in its identification, *C. auris* is becoming a critical threat to healthcare facilities globally (Jeffery-Smith et al. 2017). However, many questions remain about *C. auris*, such as how the pathogen arose and spread and how it evolved extensive multidrug resistance. A recent study demonstrated that *C. auris* clade II can acquire multidrug resistance rapidly *in vitro* (Carolus et al. 2021), providing novel insights about the mechanisms of multidrug resistance in *C. auris*. In particular, the researchers identified an ortholog of *MEC3*, which forms part of the sliding clamp for the DNA damage checkpoint (Majka and Burgers 2003), that acquired mutations during amphotericin B resistance evolution (Carolus et al. 2021). Defects in DNA damage checkpoint may result in an increasing number of A|T nucleotides in the genome, which is consistent with our observation that *C. auris* exhibits a substantial A|T mutational bias. Although the molecular mechanisms of multidrug resistance of *C. auris* require further exploration, our analysis of mutational biases provides support for the possibility that *C. auris* could acquire resistance against drugs that are cytotoxic due to oxidative DNA damage (such as amphotericin B and other polyenes) through altered DNA damage checkpoint response, which may in turn prevent apoptosis or increase mutation rate (Carolus et al. 2021; 2020; Burhans et al. 2003; Legrand et al. 2007; Healey et al. 2016).

### Species in Ascomycota vary in how their genome nucleotide compositions are affected by mutational biases and evolutionary forces

It has been previously proposed that since mutations are biased toward A|T, opposing forces such as natural selection and BGC must retain more G|C mutations to account for the variation in GC contents in species (Hershberg and Petrov 2010; Webster, Smith, and Ellegren 2003; Haddrill and Charlesworth 2008). Although these studies presented compelling evidence that gene conversion is biased toward G|C mutations in particular species and lineages (Birdsell 2002; Lesecque, Mouchiroud, and Duret 2013; Duret and Galtier 2009; Galtier et al. 2009; Pessia et al. 2012), this pattern should not be presumed to hold true for all species. For example, population genomic analysis suggests that G|C-biased gene conversion is absent in *Drosophila melanogaster* (Robinson, Stone, and Singh 2014) and analyses of mutations in tetrads from plants and fungi find no significant A|T > G|C conversion bias (Liu et al. 2018). Thus, whether BGC plays an important role in shaping genome nucleotide compositions in Ascomycota remains an open question and further research is needed to fully unravel the nature of BGC.

Even though selection has been previously shown to favor the fixation of A|T > G|C mutations in animals, plants, and other species, our data suggest that this is not always the case for fungi. The wide range of Δ_GC–GC_eq__ values in Ascomycota, from −0.04 for *S. chartarum* to 0.11 for *H. uvarum*, indicates that the selection process acting on each species varies with respect to both direction and strength (Table 2). *P. lilacinum* and *S. chartarum* both have high GC_eq_, however G|C-biased selection makes the genome of *P. lilacinum* even more GC-rich while A|T-biased selection reduces the GC content of the genome of *S. chartarum*. *C. albicans* and *C. auris* both have low GC_eq_, however A|T-biased selection makes the genome of *C. albicans* slightly less GC-rich while G|C-biased selection counteracts, to a certain extent, the loss of A|T nucleotides in the genome of *C. auris*. In addition, other species with either A|T or G|C mutational biases may not be under selection, as in the examples of *Verticillium dahliae* and *Coccidioides posadasii*. Altogether, these results demonstrate that species in Ascomycota not only vary in their mutational biases but also in how their genome nucleotide compositions are shaped by the combined effects of mutational biases and evolutionary forces, such as selection.

## Conclusion

In this study, we analyzed patterns of SNPs among 537 strains from 30 species and four classes from the fungal phylum Ascomycota. In contrary to the expectation of a universal mutational bias towards A|T, we found that species in Ascomycota demonstrate substantial diversity in their mutational biases. A predominant A|T mutational bias in Ascomycota may suggest absence or loss of DNA repair genes, although this hypothesis needs further testing. Comparison of GC equilibrium contents with observed GC contents further reveals that evolutionary forces such as selection could affect the fates of mutations by retaining either more G|C or A|T mutations. These results highlight the variation of patterns of mutations across a fungal phylum and suggest that both neutral and selective processes shape the nucleotide composition of fungal genomes.

## Acknowledgements

We thank the Rokas lab for helpful discussion and feedback. Q.Z. was partially supported by Vanderbilt Data Science Institute Summer Research Program (DSI – SRP) fellowships. J.L.S. and A.R. were supported by the Howard Hughes Medical Institute through the James H. Gilliam Fellowships for Advanced Study program. Research in A.R.’s lab is supported by grants from the National Science Foundation (DEB-1442113 and DEB-2110404), the National Institutes of Health/National Institute of Allergy and Infectious Diseases (R56 AI146096 and R01 AI153356), and the Burroughs Wellcome Fund.

## References

Akashi, H. 1995. “Inferring Weak Selection from Patterns of Polymorphism and Divergence at ‘Silent’ Sites in Drosophila DNA.” Genetics 139 (2): 1067–76. https://doi.org/10.1093/genetics/139.2.1067.

Albertin, Warren, Mathabatha E Setati, Cécile Miot-Sertier, Talitha T Mostert, Benoit Colonna-Ceccaldi, Joana Coulon, Patrick Girard, et al. 2016. “Hanseniaspora Uvarum from Winemaking Environments Show Spatial and Temporal Genetic Clustering.” Frontiers in Microbiology 6 (January): 1569. https://doi.org/10.3389/fmicb.2015.01569.

Barton, N H. 2010. “Mutation and the Evolution of Recombination.” Philosophical Transactions of the Royal Society of London. Series B, Biological Sciences 365 (1544): 1281–94. https://doi.org/10.1098/rstb.2009.0320.

Birdsell, John A. 2002. “Integrating Genomics, Bioinformatics, and Classical Genetics to Study the Effects of Recombination on Genome Evolution.” Molecular Biology and Evolution 19 (7): 1181–97. https://doi.org/10.1093/oxfordjournals.molbev.a004176.

Bont, Rinne de, and Nik van Larebeke. 2004. “Endogenous DNA Damage in Humans: A Review of Quantitative Data.” Mutagenesis 19 (3): 169–85. https://doi.org/10.1093/mutage/geh025.

Budden, Timothy, and Nikola A Bowden. 2013. “The Role of Altered Nucleotide Excision Repair and UVB-Induced DNA Damage in Melanomagenesis.” International Journal of Molecular Sciences 14 (1): 1132–51. https://doi.org/10.3390/ijms14011132.

Burhans, William C, Martin Weinberger, Maria A Marchetti, Lakshmi Ramachandran, Gennaro D’Urso, and Joel A Huberman. 2003. “Apoptosis-like Yeast Cell Death in Response to DNA Damage and Replication Defects.” Mutation Research/Fundamental and Molecular Mechanisms of Mutagenesis 532 (1): 227–43. https://doi.org/https://doi.org/10.1016/j.mrfmmm.2003.08.019.

Cambareri, E B, B C Jensen, E Schabtach, and E U Selker. 1989. “Repeat-Induced G-C to A-T Mutations in Neurospora.” Article. Science (American Association for the Advancement of Science) 244 (4912): 1571–75. https://doi.org/10.1126/science.2544994.

Carolus, Hans, Siebe Pierson, Katrien Lagrou, and Patrick van Dijck. 2020. “Amphotericin B and Other Polyenes-Discovery, Clinical Use, Mode of Action and Drug Resistance.” Journal of Fungi (Basel, Switzerland) 6 (4): 321. https://doi.org/10.3390/jof6040321.

Carolus, Hans, Siebe Pierson, José F Muñoz, Ana Subotić, Rita B Cruz, Christina A Cuomo, and Patrick van Dijck. 2021. “Genome-Wide Analysis of Experimentally Evolved Candida Auris Reveals Multiple Novel Mechanisms of Multidrug Resistance.” MBio 12 (2): e03333–20. https://doi.org/10.1128/mBio.03333-20.

Duret, Laurent, and Nicolas Galtier. 2009. “Biased Gene Conversion and the Evolution of Mammalian Genomic Landscapes.” Annual Review of Genomics and Human Genetics 10 (1): 285–311. https://doi.org/10.1146/annurev-genom-082908-150001.

Eyre-Walker, Adam, and Peter D Keightley. 2007. “The Distribution of Fitness Effects of New Mutations.” Nature Reviews Genetics 8 (8): 610–18. https://doi.org/10.1038/nrg2146.

Galagan, James E, and Eric U Selker. 2004. “RIP: The Evolutionary Cost of Genome Defense.” Article. Trends in Genetics 20 (9): 417–23. https://doi.org/10.1016/j.tig.2004.07.007.

Galtier, Nicolas, Laurent Duret, Sylvain Glémin, and Vincent Ranwez. 2009. “GC-Biased Gene Conversion Promotes the Fixation of Deleterious Amino Acid Changes in Primates.” Trends in Genetics 25 (1): 1–5. https://doi.org/https://doi.org/10.1016/j.tig.2008.10.011.

Haag-Liautard, Cathy, Mark Dorris, Xulio Maside, Steven Macaskill, Daniel L Halligan, Brian Charlesworth, and Peter D Keightley. 2007. “Direct Estimation of per Nucleotide and Genomic Deleterious Mutation Rates in Drosophila.” Nature 445 (7123): 82–85. https://doi.org/10.1038/nature05388.

Haddrill, Penelope R, and Brian Charlesworth. 2008. “Non-Neutral Processes Drive the Nucleotide Composition of Non-Coding Sequences in Drosophila.” Biology Letters 4 (4): 438–41. https://doi.org/10.1098/rsbl.2008.0174.

Hane, James K, Angela H Williams, Adam P Taranto, Peter S Solomon, and Richard P Oliver. 2014. “Repeat-Induced Point Mutation: A Fungal-Specific, Endogenous Mutagenesis Process.” Bookitem. In Genetic Transformation Systems in Fungi, Volume 12, 55–68. Fungal Biology. Cham: Springer International Publishing. https://doi.org/10.1007/978-3-319-10503-1_4.

Hawksworth, David, and Robert Lücking. 2017. “Fungal Diversity Revisited: 2.2 to 3.8 Million Species.” Microbiology Spectrum 5 (July). https://doi.org/10.1128/microbiolspec.FUNK-0052-2016.

Healey, Kelley R, Cristina Jimenez Ortigosa, Erika Shor, and David S Perlin. 2016. “Genetic Drivers of Multidrug Resistance in Candida Glabrata.” Frontiers in Microbiology 7 (December): 1995. https://doi.org/10.3389/fmicb.2016.01995.

Hershberg, Ruth, and Dmitri A Petrov. 2010. “Evidence That Mutation Is Universally Biased towards AT in Bacteria.” PLOS Genetics 6 (9): e1001115-. https://doi.org/10.1371/journal.pgen.1001115.

Hierro, Núria, Ángel González, Albert Mas, and Jose M Guillamón. 2006. “Diversity and Evolution of Non-Saccharomyces Yeast Populations during Wine Fermentation: Effect of Grape Ripeness and Cold Maceration.” FEMS Yeast Research 6 (1): 102–11. https://doi.org/https://doi.org/10.1111/j.1567-1364.2005.00014.x.

Horns, Felix, Elsa Petit, Roxana Yockteng, and Michael E Hood. 2012. “Patterns of Repeat-Induced Point Mutation in Transposable Elements of Basidiomycete Fungi.” Genome Biology and Evolution 4 (3): 240–47. https://doi.org/10.1093/gbe/evs005.

James, Timothy Y, Jason E Stajich, Chris Todd Hittinger, and Antonis Rokas. 2020. “Toward a Fully Resolved Fungal Tree of Life.” Annual Review of Microbiology 74 (1): 291–313. https://doi.org/10.1146/annurev-micro-022020-051835.

Jeffery-Smith, Anna, Surabhi K Taori, Silke Schelenz, Katie Jeffery, Elizabeth M Johnson, Andrew Borman, Candida auris Incident Management Team, Rohini Manuel, and Colin S Brown. 2017. “Candida Auris: A Review of the Literature.” Clinical Microbiology Reviews 31 (1): e00029–17. https://doi.org/10.1128/CMR.00029-17.

Jos, Houbraken, Visagie Cobus M, Frisvad Jens C, and Rokas Antonis. 2022. “Recommendations To Prevent Taxonomic Misidentification of Genome-Sequenced Fungal Strains.” Microbiology Resource Announcements 10 (48): e01074–20. https://doi.org/10.1128/MRA.01074-20.

Keightley, Peter D, Urmi Trivedi, Marian Thomson, Fiona Oliver, Sujai Kumar, and Mark L Blaxter. 2009. “Analysis of the Genome Sequences of Three Drosophila Melanogaster Spontaneous Mutation Accumulation Lines.” Genome Research 19 (7): 1195–1201. https://doi.org/10.1101/gr.091231.109.

Kurtz, Stefan, Adam Phillippy, Arthur L Delcher, Michael Smoot, Martin Shumway, Corina Antonescu, and Steven L Salzberg. 2004. “Versatile and Open Software for Comparing Large Genomes.” Genome Biology 5 (2): R12. https://doi.org/10.1186/gb-2004-5-2-r12.

Legrand, Melanie, Christine L Chan, Peter A Jauert, and David T Kirkpatrick. 2007. “Role of DNA Mismatch Repair and Double-Strand Break Repair in Genome Stability and Antifungal Drug Resistance in Candida Albicans.” Eukaryotic Cell 6 (12): 2194–2205. https://doi.org/10.1128/EC.00299-07.

Lesecque, Yann, Dominique Mouchiroud, and Laurent Duret. 2013. “GC-Biased Gene Conversion in Yeast Is Specifically Associated with Crossovers: Molecular Mechanisms and Evolutionary Significance.” Molecular Biology and Evolution 30 (6): 1409–19. https://doi.org/10.1093/molbev/mst056.

Li, Yuanning, Jacob L Steenwyk, Ying Chang, Yan Wang, Timothy Y James, Jason E Stajich, Joseph W Spatafora, et al. 2021. “A Genome-Scale Phylogeny of the Kingdom Fungi.” Current Biology 31 (8): 1653–1665.e5. https://doi.org/10.1016/j.cub.2021.01.074.

Liu, Haoxuan, Ju Huang, Xiaoguang Sun, Jing Li, Yingwen Hu, Luyao Yu, Gianni Liti, Dacheng Tian, Laurence D Hurst, and Sihai Yang. 2018. “Tetrad Analysis in Plants and Fungi Finds Large Differences in Gene Conversion Rates but No GC Bias.” Nature Ecology & Evolution 2 (1): 164–73. https://doi.org/10.1038/s41559-017-0372-7.

Lynch, Michael. 2010. “Rate, Molecular Spectrum, and Consequences of Human Mutation.” Proceedings of the National Academy of Sciences 107 (3): 961. https://doi.org/10.1073/pnas.0912629107.

Majka, Jerzy, and Peter M J Burgers. 2003. “Yeast Rad17/Mec3/Ddc1: A Sliding Clamp for the DNA Damage Checkpoint.” Proceedings of the National Academy of Sciences of the United States of America 100 (5): 2249–54. https://doi.org/10.1073/pnas.0437148100.

Messer, Philipp W. 2009. “Measuring the Rates of Spontaneous Mutation From Deep and Large-Scale Polymorphism Data.” Genetics 182 (4): 1219. https://doi.org/10.1534/genetics.109.105692.

Monroe, J Grey, Thanvi Srikant, Pablo Carbonell-Bejerano, Claude Becker, Mariele Lensink, Moises Exposito-Alonso, Marie Klein, et al. 2022. “Mutation Bias Reflects Natural Selection in Arabidopsis Thaliana.” Nature. https://doi.org/10.1038/s41586-021-04269-6.

Naranjo-Ortiz, Miguel A, and Toni Gabaldón. 2019. “Fungal Evolution: Diversity, Taxonomy and Phylogeny of the Fungi.” Biological Reviews of the Cambridge Philosophical Society 94 (6): 2101–37. https://doi.org/10.1111/brv.12550.

Orr, H Allen. 2005. “The Genetic Theory of Adaptation: A Brief History.” Nature Reviews Genetics 6 (2): 119–27. https://doi.org/10.1038/nrg1523.

Pessia, Eugénie, Alexandra Popa, Sylvain Mousset, Clément Rezvoy, Laurent Duret, and Gabriel A B Marais. 2012. “Evidence for Widespread GC-Biased Gene Conversion in Eukaryotes.” Genome Biology and Evolution 4 (7): 675–82. https://doi.org/10.1093/gbe/evs052.

Robinson, Matthew C, Eric A Stone, and Nadia D Singh. 2014. “Population Genomic Analysis Reveals No Evidence for GC-Biased Gene Conversion in Drosophila Melanogaster.” Molecular Biology and Evolution 31 (2): 425–33. https://doi.org/10.1093/molbev/mst220.

Satoh, Kazuo, Koichi Makimura, Yayoi Hasumi, Yayoi Nishiyama, Katsuhisa Uchida, and Hideyo Yamaguchi. 2009. “Candida Auris Sp. Nov., a Novel Ascomycetous Yeast Isolated from the External Ear Canal of an Inpatient in a Japanese Hospital.” Microbiology and Immunology 53 (1): 41–44. https://doi.org/https://doi.org/10.1111/j.1348-0421.2008.00083.x.

Schütz, M, and J Gafner. 2008. “Analysis of Yeast Diversity during Spontaneous and Induced Alcoholic Fermentations.” Journal of Applied Microbiology 75 (March): 551–58. https://doi.org/10.1111/j.1365-2672.1993.tb01594.x.

Selker, Eric U, Edward B Cambareri, Bryan C Jensen, and Kenneth R Haack. 1987. “Rearrangement of Duplicated DNA in Specialized Cells of Neurospora.” Article. Cell 51 (5): 741–52. https://doi.org/10.1016/0092-8674(87)90097-3.

Shen, Xing-Xing, Jacob L Steenwyk, Abigail L LaBella, Dana A Opulente, Xiaofan Zhou, Jacek Kominek, Yuanning Li, Marizeth Groenewald, Chris T Hittinger, and Antonis Rokas. 2020. “Genome-Scale Phylogeny and Contrasting Modes of Genome Evolution in the Fungal Phylum Ascomycota.” Science Advances 6 (45): eabd0079. https://doi.org/10.1126/sciadv.abd0079.

Steenwyk, Jacob L, Dana A Opulente, Jacek Kominek, Xing-Xing Shen, Xiaofan Zhou, Abigail L Labella, Noah P Bradley, et al. 2019. “Extensive Loss of Cell-Cycle and DNA Repair Genes in an Ancient Lineage of Bipolar Budding Yeasts.” PLOS Biology 17 (5): e3000255-. https://doi.org/10.1371/journal.pbio.3000255.

Webster, Matthew T, Nick G C Smith, and Hans Ellegren. 2003. “Compositional Evolution of Noncoding DNA in the Human and Chimpanzee Genomes.” Molecular Biology and Evolution 20 (2): 278–86. https://doi.org/10.1093/molbev/msg037.

Zhu, Yuan O, Mark L Siegal, David W Hall, and Dmitri A Petrov. 2014. “Precise Estimates of Mutation Rate and Spectrum in Yeast.” Proceedings of the National Academy of Sciences 111 (22): E2310. https://doi.org/10.1073/pnas.1323011111.

