## Supplementary Figures for "Lack of universal mutational biases in a fungal phylum"

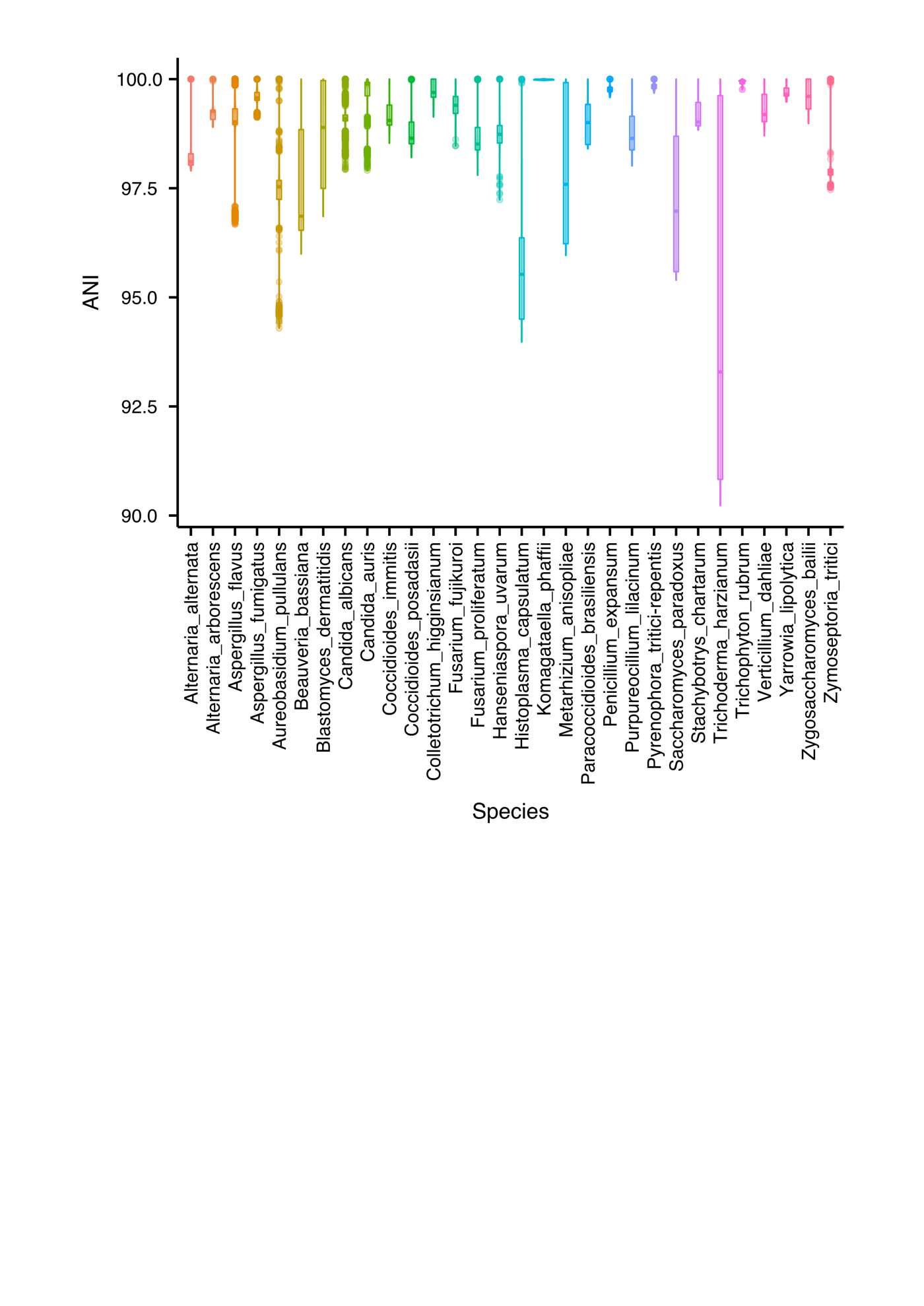

**Figure S1. Average nucleotide identity between the genomes of strains within each of the 30 Ascomycota species.**

The average nucleotide identity (ANI) is calculated using the FastANI program. Distribution of intraspecies ANI values for each species is shown as one boxplot. Strains with intraspecies ANI values below 90 and obvious outliers were removed from our study (prior to this analysis). All species except *Trichoderma harzianum* have median intraspecies ANI values greater than 95. *Trichoderma harzianum* has the lowest intraspecies ANI value, though still above 90.

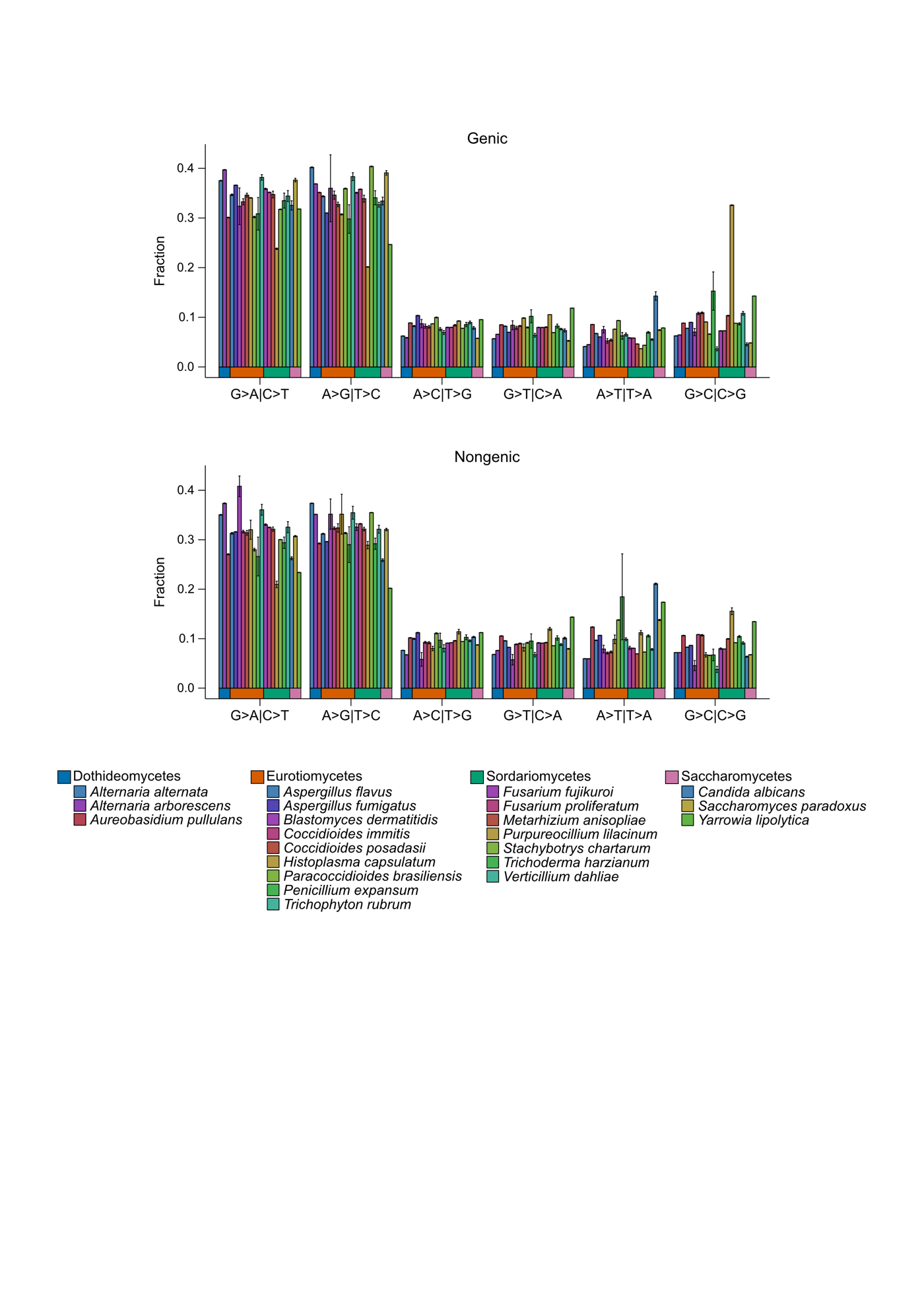

**Figure S2. Relative genome-wide rates of protein-coding and non-coding nucleotide substitutions.**

The most common types of substitutions observed are G>A | C>T and A>G | T>C transitions in both protein-coding and non-coding regions. The overall mutational pattern is similar across protein-coding and non-coding regions. Note that eight species were not included in these analyses as they currently lack publicly available genome annotations.

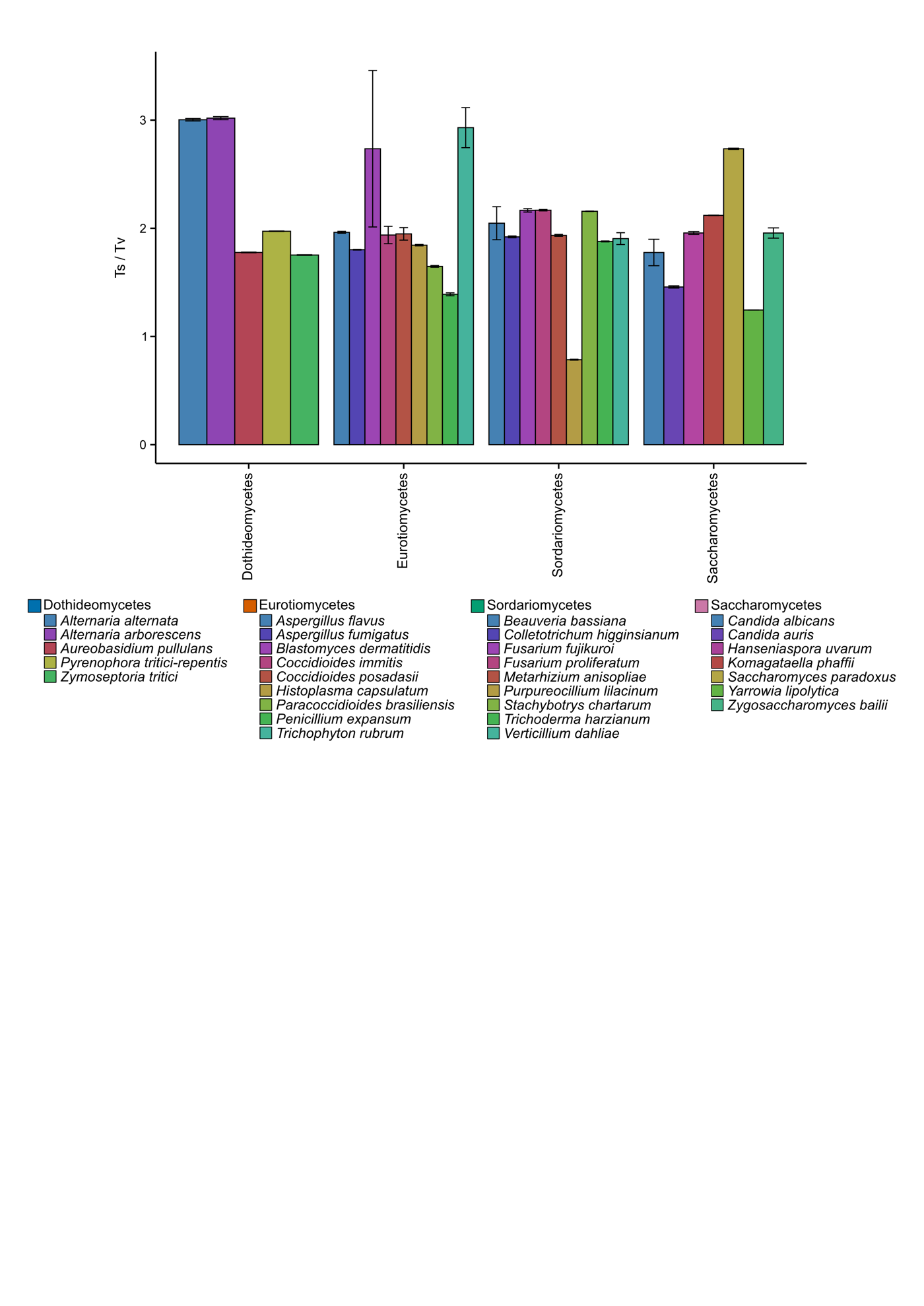

**Figure S3. Transition / transversion ratios among Ascomycota fungi.**

The transition / transversion (Ts/Tv) ratios reveal most species have Ts/Tv ratios around two, indicating that transitions occur more frequently than transversions as expected. One notable exception is *Purpureocillium lilacinum* (Sordariomycetes), whose Ts/Tv ratio is below one.

**Supplementary Tables**

**Table S1. Statistics of the genomes of all strains used in this study.**
